# Pairwise library screen systematically interrogates *Staphylococcus aureus* Cas9 specificity in human cells

**DOI:** 10.1101/269399

**Authors:** Josh Tycko, Luis A. Barrera, Nicholas Huston, Ari E. Friedland, Xuebing Wu, Jonathan S. Gootenberg, Omar O. Abudayyeh, Vic E. Myer, Christopher J. Wilson, Patrick D. Hsu

**Affiliations:** Editas Medicine, 11 Hurley St., Cambridge, MA 02141, USA; Department of Molecular Biophysics and Biochemistry, Yale University, New Haven, CT 06511, USA; Whitehead Institute for Biomedical Research, Cambridge, MA 02142, USA; Department of Systems Biology, Harvard, Cambridge, MA 02138, USA; Department of Health Sciences and Technology, Massachusetts Institute of Technology, Cambridge, MA 02139, USA

## Abstract

We report a high-throughput screening approach to measure *Staphylococcus aureus* Cas9 (SaCas9) genome editing variation in human cells across a large repertoire of 88,692 single guide RNAs (sgRNAs) paired with matched or mismatched target sites in a synthetic cassette. We incorporated randomized barcodes that enable ‘whitelisting’ of correctly synthesized molecules for further downstream analysis, in order to circumvent the limitation of oligonucleotide synthesis errors. We find SaCas9 sgRNAs with a 21-nucleotide spacer are most active against off-targets with single and double mismatches, compared to shorter or longer sgRNAs. Using this dataset, we developed an SaCas9 specificity model that performs well in ranking off-target sites. The barcoded pairwise library screen enabled high-fidelity recovery of guide-target relationships, providing a scalable framework for the investigation of CRISPR enzyme properties and general nucleic acid interactions.

## Main text

The compact SaCas9 enables *in vivo* delivery of Cas9 and multiple sgRNA packaged within a single adeno-associated virus (AAV) vector^1,2^, serving as a promising platform for gene editing therapies. AAV-SaCas9 is capable of achieving therapeutic levels of genome editing in preclinical animal models of Duchenne muscular dystrophy^3,4^, ornithine transcarbamylase deficiency^5^, and of HIV infection^6^. Translating these promising initial results into medicines requires a rigorous understanding of intended and unintended genome editing. SaCas9-mediated off-target effects have been detected with genome-wide methods, including GUIDE-seq^7^ and BLESS^1,8^, and direct visualization of dSaCas9-EGFP binding in cells^9^. However, the sequence determinants of SaCas9 cleavage specificity have not been profiled. Furthermore, SaCas9 is known to efficiently cleave genomic DNA with spacer lengths from 20 – 24 nt^1,2^, but the effect of spacer length on specificity is not known.

To systematically investigate SaCas9 specificity in human cells, we developed a method to test a library of sgRNAs against a library of genome-integrated synthetic target sequences. Lentiviral delivery of the ‘pairwise library’ cassette results in integration of a sgRNA and paired synthetic target site in the genome (**Fig. 1A**). We first designed a library of 88,692 sgRNA-target pairs, distributed among 73 sgRNA groups (**Table 1**). Within a group, the sgRNA had shared sequence in positions 1-18 and ranged in length from 19 nt to 24 nt spacers. All sgRNAs were paired with target sites bearing all possible single mismatches and subsets of sgRNAs were paired with all possible double mismatches or all possible double transversions. Five groups of sgRNA were paired with target sites bearing all possible single insertions and deletions, to study the effect of DNA and RNA bulges. The protospacer adjacent motif (PAM) was held constant at 5’-CAGGGT-3’ to match the consensus sequence of 5’-NNGRRT-3’ ^1^.

**Figure 1.**
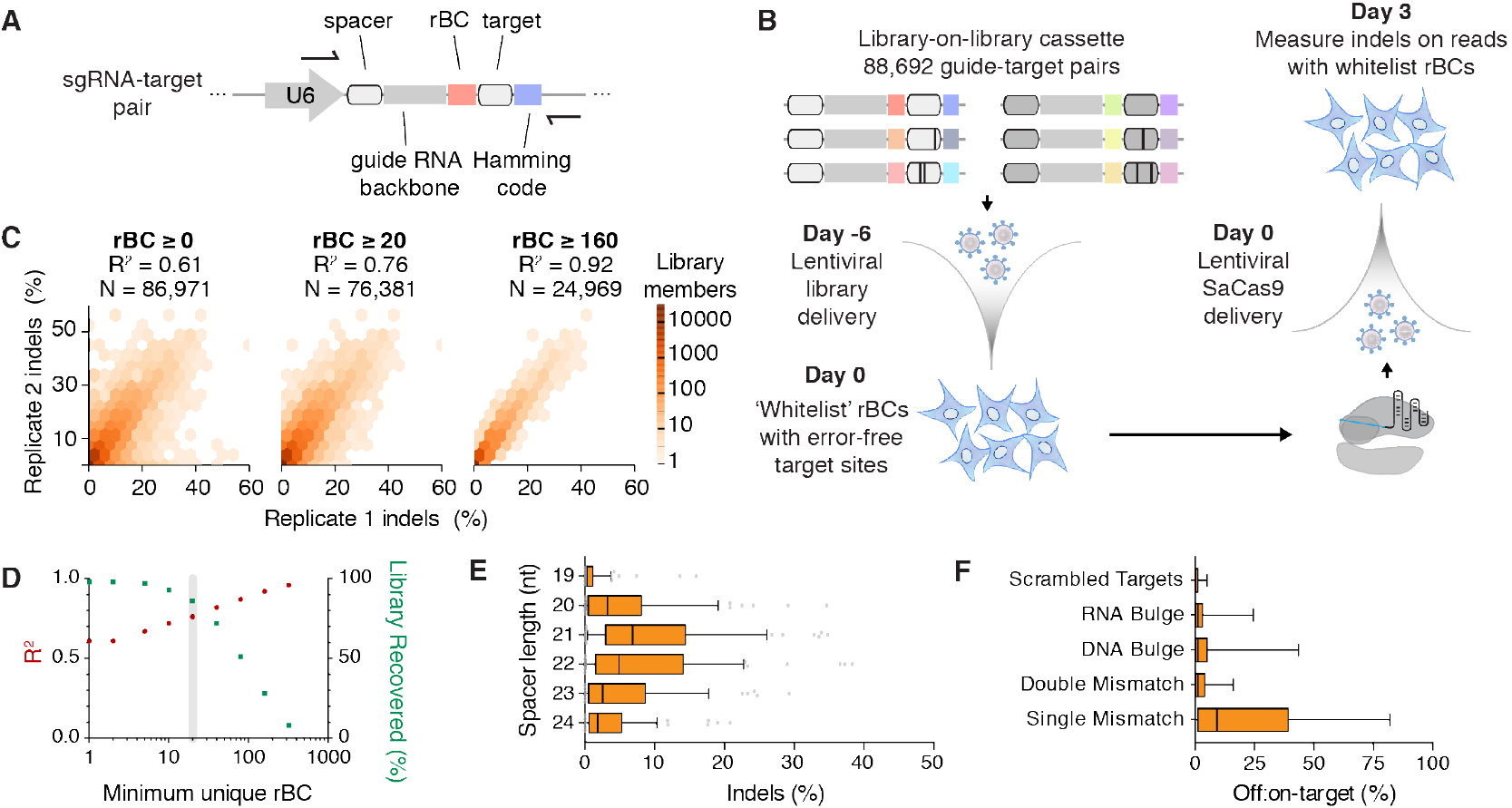
Pairwise library screen of SaCas9 genome editing specificity. **A.** Schematic of the pairwise library cassette. Individual library members have variable spacer and target sequences, and each member is identifiable by a unique 15 nt error-correcting Hamming barcode. Individual molecules of each library member are tagged with a unique randomized barcode (rBC). **B.** Schematic of the pooled pairwise library screen workflow. Each sgRNA-target pair is associated with many rBCs. The library was initially installed in HEK 293T cells by lentivirus and sequenced to generate a whitelist of rBCs associated with error-free guide-target pairs. SaCas9 was then delivered by lentivirus. After 3 days, the library was sequenced to measure Cas9-mediated indels. **C.** Reproducibility of a pairwise library screen increases if a greater number of whitelist rBCs is required for each library member. Hexagon heat color represents the density of library members, while white area represents 0 library members. **D.** The fraction of recovered library members decreases as a greater number of rBCs is required. All downstream analyses were performed with a minimum of 20 unique whitelist rBCs for each library member (grey). **E.** On-target indel efficiency for SaCas9 guide-target pairs, binned by spacer length. Boxes denote median and interquartile range (IQR) while whiskers extend to the 10^th^ and 90^th^ percentile (n = 653 sgRNA-target pairs). **F.** Comparison of SaCas9 activity across categories of target sites. The ‘scrambled targets’ are negative controls. Boxes denote median and IQR while whiskers extend to the 10^th^ and 90^th^ percentile (n = 88,692 sgRNA-target pairs).

We next measured the genome editing activity of these guide-target pairs in human cells (**Fig. 1B**). The library cassette lentivirus was transduced in HEK 293FT cells at low multiplicity-of-infection (MOI) to enrich for single-copy integration events, ensuring independent editing reactions per cell. Genomic DNA was extracted 0 and 3 days after SaCas9 transduction and the library cassette was PCR-amplified prior to Illumina sequencing.

After editing has occurred, insertions or deletions (indels) in the target site can obscure the original sgRNA-target pair relationship. Accordingly, each library member (i.e., a unique sgRNA-target pair) was linked with a unique, error-correcting Hamming barcode^10^ downstream of the targeted PAM. Post-editing, this Hamming barcode identifies the original sgRNA-target sequence in a sequence read. The indel frequencies associated with each Hamming barcode determine editing efficiency for each sgRNA-target pair.

Reasoning that a subset of molecules representing a particular library member would be subject to synthesis errors that generate an inappropriate mismatch or indel, we appended an additional randomized barcode (rBC) downstream of the sgRNA. These rBC effectively barcode unique lentiviral integrations. Sequencing from Day 0 was used to ‘whitelist’ the guide-target cassettes that were error-free and sufficiently represented prior to Cas9 delivery. This whitelist minimizes false positive indels that arise from synthesis errors or other causes, and improves the reproducibility of pairwise library screens by filtering out library members with insufficient cellular representation (i.e., library members with a low number of rBCs).

Indel rates and the corresponding off:on-target ratios at Day 3 were then computed using only whitelisted rBCs (i.e., cassettes that were error-free at Day 0). Indel levels were observed to be increasingly reproducible as the minimum number of unique whitelist rBCs per library member was also increased (**Fig. 1C**). After filtering for library members with at least 20 independent integrations (ie. ≥ 20 rBCs), 83% of the library remained and the biological replicates were correlated (R^2^ = 0.76) (**Fig. 1D**).

We first analyzed the on-target activity of SaCas9. Based on our observation of saturation in indel levels by Day 6 in a pilot time-course study (**Fig. S1**), we sequenced the screen samples on Day 3 to determine representative but non-saturated indel efficiency. 21 nt and 22 nt spacers edited on-target sites most efficiently, with no significant difference between on-target activity distributions with these two lengths (**Fig. 1E**). However, 21 nt spacers (but not 22 nt spacers) were significantly more efficient than 23 nt spacers (*p*<0.05, Dunn’s multiple comparisons test). This result refines smaller-scale studies (of 4 and 21 sgRNA) that had defined the optimal spacer length as being 21 - 23 nt ^1^ or 20 - 24 nt ^2^.

Next, we compared the average off-target indel rates for mismatched sgRNA-target library members and found targets with single mismatches were edited 3.4-fold less than matched targets on average (p<0.01, Dunn’s multiple comparisons test) (**Fig. 1F**), while the targets with double mismatches or bulges were edited even less frequently.

SaCas9 off:on-target activity can be further considered as a function of both mismatch identity and position. Overall, mismatch tolerance was low in the ~9 nt PAM-proximal ‘seed’ region, oscillated between positions 13 – 19, and was higher at the PAM-distal region (**Fig. 2A, B**). This tolerance pattern was consistent across spacer lengths, but the 21 nt spacer was the most tolerant of mismatches at any position.

**Figure 2.**
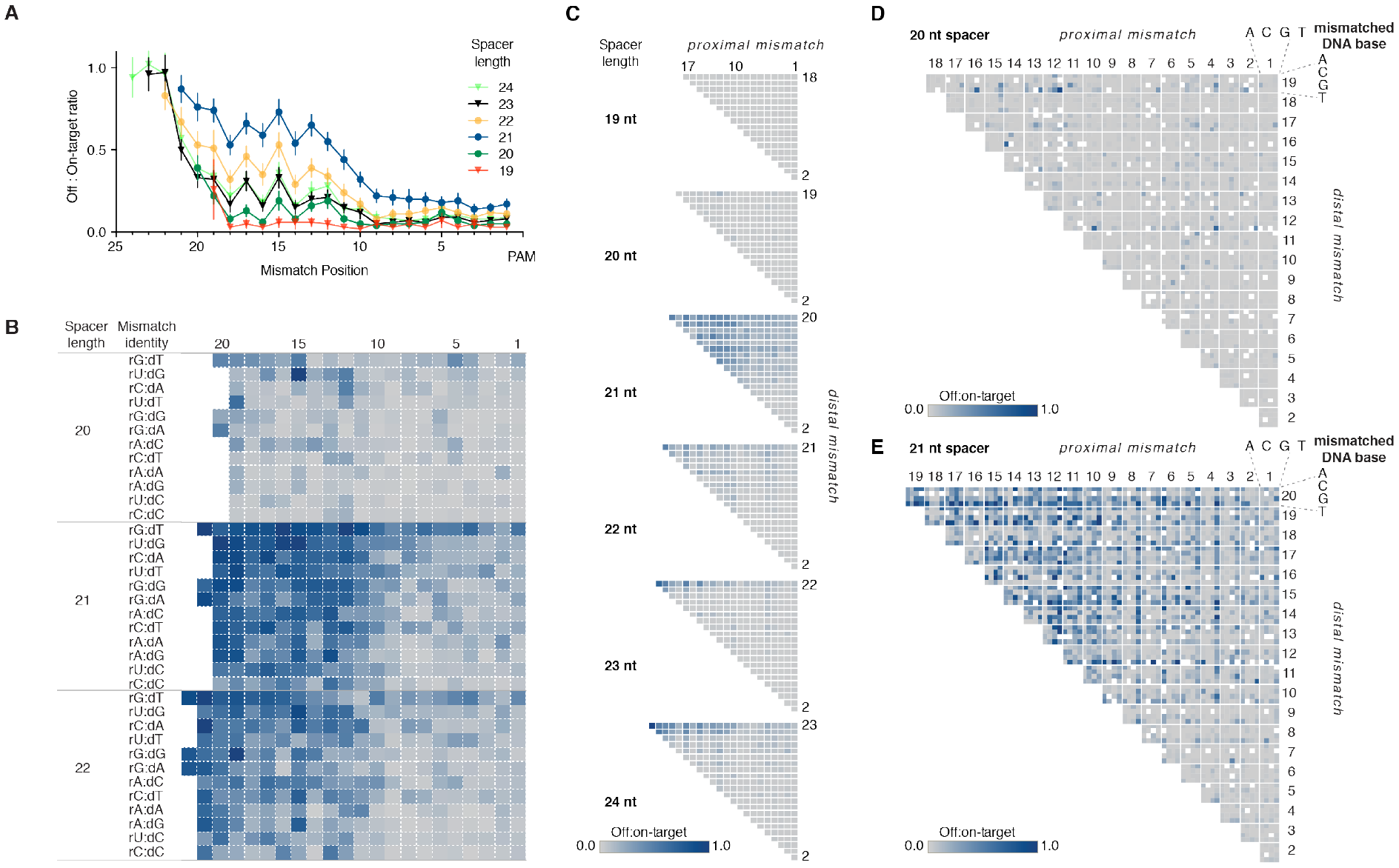
21 nt spacers are most tolerant of single and double mismatches. **A.** Average effect of sgRNA spacer length and mismatch position on SaCas9 single mismatch tolerance. Mean ± 95% confidence interval is shown (n = 16,742 sgRNA-target pairs). **B.** Heatmap of relative SaCas9 cleavage efficiency for each possible RNA:DNA base pair, calculated for all single mismatch library members. (n = 10,344 sgRNA-target pairs) **C.** Heatmap of relative SaCas9 double mismatch tolerance across different spacer lengths and positions. Data is aggregated from 10 sgRNA for which all possible single and double mismatches were tested (n = 50,053 sgRNA-target pairs). **D.** DNA base identity and position effect on double mismatch tolerance for 20 nt spacer sgRNA. **E.** DNA base identity and position effect on double mismatch tolerance for 21 nt spacer sgRNA.

21 nt spacers were also more tolerant of double mismatches than any other spacer length (*p*<0.0001, Dunn’s multiple comparisons test), in a position-dependent manner (**Fig. 2C, E**). 21 nt spacers had a mean 16% off:on target activity ratio while we observed a mean of only 2% and 6% off:on-target activity across all double-mismatched sites targeted by 20 nt or 22 nt spacers, respectively (**Fig. 2C, D**). Upon inspecting the subset of library members with insertions or deletions in the target site, we found that both RNA and DNA bulges in the sgRNA:target duplex were minimally tolerated at positions 1-18, regardless of sgRNA length (**Fig. S2**). However, PAM-distal bulges were near-completely tolerated and the bulge nucleotide identity had no significant effect on the indel rate (**Fig. S2**).

Given the striking effect of the spacer length on specificity, we conducted Northern blot analysis to confirm that this range of spacer lengths is reliably maintained in human cells. sgRNA from 18- to 24 nt were accurately maintained at full length when expressed from a U6 promoter, regardless of whether or not the 5’ nucleotide was a guanine (**Fig. S3**). Further, sgRNA were hardly detectable in cells lacking SaCas9 expression.

To validate these findings with an orthogonal method, we designed GFP-targeting sgRNAs with variable length spacers and synthesized sgRNA with single mismatches at every position. In independent wells, we transiently transfected SaCas9 and each sgRNA into a stable HEK 293T-GFP cell line and measured GFP knockout by flow cytometry. Consistent with the screen, we observed low tolerance of PAM-proximal mismatches, and more variable PAM-distal specificity in a mismatch identity and nucleotide position-dependent manner. As in the screen, we again found 21 nt spacers to be most tolerant of single mismatches (**Fig. S4**).

We next formulated a non-linear regression specificity score, trained from our screen dataset, to integrate the relative contribution of mismatch position, identity, number, and density. These scores correlate well with the observed off-target activity in the screen (**Fig. S5**). Our model assumes that mismatches can be modeled independently, and performs similarly on off-targets with single or multiple mismatches (**Fig. S5**). This scoring algorithm was capable of predicting SaCas9 specificity in the orthogonal GFP mismatch assay with high performance (**Fig. 3A**). Furthermore, the score performed well in ranking endogenous off-target sites, as measured by GUIDE-seq^7,11^, with a Spearman correlation of 0.82 (**Fig. 3B**).

**Figure 3.**
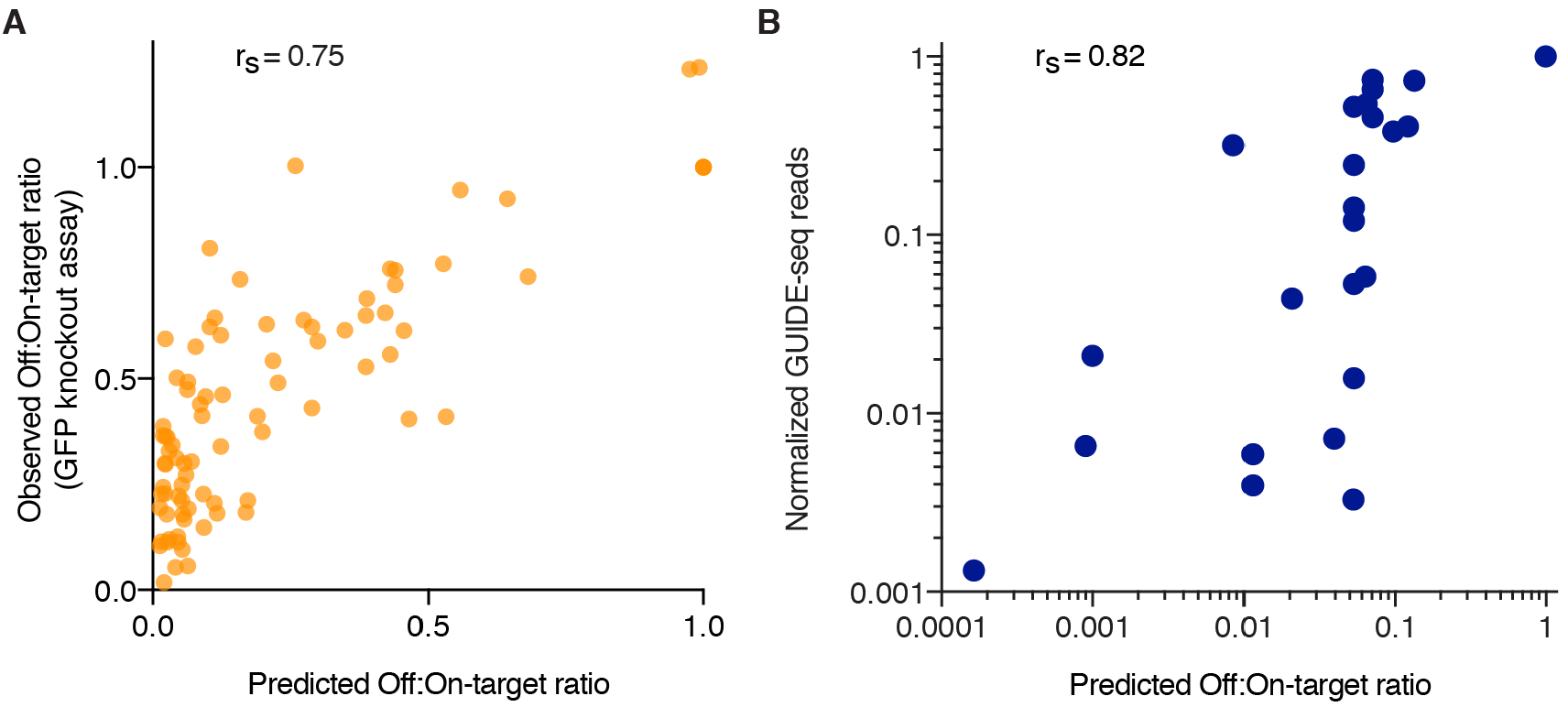
SaCas9 specificity model. **A.** A regression model trained on the pairwise library screen data correlates with an independent validation set (n = 86 sgRNA-target pairs). The sgRNAs knockout a stable GFP and include single mismatches in the spacers to mimic off-target effects. **B.** Spearman rank correlation of the SaCas9 specificity model compared with previously reported SaCas9 GUIDE-seq data^7^.

## Discussion

These findings support a simple strategy to mitigate the risk of off-target activity by adjusting the spacer length. Most often, 22 nt spacers offer the optimal balance of efficiency and specificity, while 20 nt spacers improve specificity at the cost of lesser efficiency. This data also supports known strategies such as selecting sgRNA with maximal sequence dissimilarity from off-target sites, and particularly avoiding off-target sites with only PAM-distal mismatches. Finally, our specificity model trained on SaCas9-specific parameters can be used for the *in silico* selection of guides and to prioritize off-target sites for follow-up.

The challenge of characterizing genome editing and nucleic acid specificity is well-suited to high-throughput approaches because of the large space of possible guide-target pairs. While genome-wide off-target detection methods^1,11–16^ are important for characterizing individual sgRNA of interest, they are also limited by the availability of endogenous off-target sites, and do not provide general models of specificity. A complementary approach is to generate a synthetic library that more thoroughly covers the space of possible off-target sites. To date, such studies have primarily been performed *in vitro*^17,18^ and in yeast^19^. While specificity of the Cas9 from *S. pyogenes* has been previously profiled in human cells via cell surface marker knockout and flow cytometry^20^, the pairwise library approach described here provides an important alternative method. Previous reports suggest that even single-nucleotide changes in a sgRNA can strongly affect activity by changing RNA secondary structure^21^, but our screen avoids this concern and mimics a real-world scenario by placing the mismatches in the target DNA. Further, the edits are directly measured at the DNA sequence. This property could additionally enable high-throughput evaluation of nuclease-mediated DNA repair outcomes in future studies.

A related approach was recently employed to characterize Cpf1 as a genome editing tool^22^ by screening linked libraries, but was sometimes limited by high error rates at baseline. In one experiment, only 2% of their library had sufficient baseline reads without indels to qualify for further analysis. In contrast, our design includes two barcodes, one for the target-guide pair and another to track each integration event, which allows us to whitelist the targets that were error-free at baseline and recover the majority of the library for analysis. Low MOI delivery of our pairwise library further facilitates the measurement of independent editing reactions for each sgRNA-target pair compartmentalized in individual cells.

Our barcoded pairwise library screening approach provides a general framework for understanding and engineering nucleic acid interactions, and could be exploited for oligonucleotide probe or switch design. We demonstrate its utility via high-throughput characterization of SaCas9 specificity, which could be extended to interrogate other nuclease-centric properties such as DNA repair outcomes in therapeutically relevant cell types.

## Materials and Methods

### Library design and cloning

Custom Python scripts were written for the pairwise library design. Initially, we generated a random library of 19-24 nt spacer sequences. To minimize undesired Cas9 targeting outside the lentivirally-integrated pairwise library cassette, the sgRNA sequences were then computationally optimized to be highly orthogonal to the human reference genome by filtering the list of candidate spacers against the hg19 assembly. A 5’ G was held constant for every unique sgRNA spacer for reliable U6-driven expression. The (PAM) was held constant at 5’-CAGGGT-3’. The number of single-mismatch, double-mismatch, DNA bulge, RNA bulge, and control guides that were generated for each unique sgRNA spacer is described in **Table S1**.

Hamming codes were generated using a modified version of a Python script available on Github (https://github.com/mdshw5/hamstring). We modified the code to increase the barcode length to 15-bp, expanding the number of available barcodes so as to cover the whole library. These barcodes are composed of 10 data bases, 4 checksum bases, and 1 parity base. We excluded homopolymers of >3 nt and filtered for GC content from 30% to 70%. This resulted in 812,547 barcodes, which were sub-sampled to 93,000, enabling an increased edit-distance greater than 2.

The library was synthesized as a pooled library of 88,692 single stranded 135-142 nt oligonucleotides by CustomArray. Variable oligo length is due to the varied spacer lengths. In order to accommodate constraints of synthesis length, oligo synthesis did not include the full length sgRNA tail, but instead included a short BsmBI Type IIS “tracrRNA cloning site” in between the spacer and target sequences.

First, the library oligo was PCR amplified and cloned by Gibson Assembly (NEB) into the pairwise library lentiviral backbone, which included a U6 promoter for sgRNA expression and a puromycin resistance gene. The plasmid was then electroporated into Endura ElectroCompetent cells (Biorad). To maintain library complexity, the transformed cells were plated on large 245 × 245 mm LB plates (Teknova) and colony density was estimated by serially diluting and spreading the transformed cells on LB plates. Colonies were quantified with an online image analysis tool (Benchling).

This “pre-tracrRNA” pairwise library was sequenced by MiSeq to verify its representation and rate of synthesis errors. The tracrRNA was synthesized as a PAGE-purified ultramer (Integrated DNA Technologies), with an additional 8 nt 5’- NNNNNNNN-3’ randomized barcode (rBC) at the 3’ end to identify independent DNA molecules representing each library member. The tracrRNA-rBC oligo was PCR amplified and ligated into the BsmBI cloning site of the pre-tracrRNA. The resulting plasmid pool was also subjected to deep sequencing by MiSeq to verify library member representation and synthesis quality.

### Cell Culture

HEK293T and HEK293FT (Life Technologies, catalog #R700-07) cells were cultured in Dulbecco’s modified eagle medium (DMEM), supplemented with 10% fetal bovine serum (FBS) and 1% penicillin-streptomyocin (D10). Cells were maintained in T225 flasks while screening. The antibiotic selections used 10 *μ*g/ml blasticidin or 0.5 *μ*g/ml puromycin. HEK293-GFP (GenTarget, catalog #SC001) cells were maintained in DMEM, supplemented with 10% FBS, 5% penicillin-streptomycin, and 2 mM Glutamax. All HEK293 were kept at 37°C in a 5% CO_2_ incubator.

### Lentiviral production and titering

To package lentivirus, the library plasmid was co-transfected with pMD2.G and psPAX2 into 120 million HEK293FT cells, using Lipofectamine 2000 (Thermo Fisher) in 10 T-225 flasks (Sigma). After 72 hours, the supernatant was harvested and concentrated according to the LentiX Concentrator protocol, then stored at −80°C. In order to titer this lentiviral preparation, we spinfected 3 million HEK293T cells per well with 0, 40, 80, 120, 160, or 200 *μ*l of the lentiviral supernatant in 2 ml D10 media supplemented with 8 *μ*g/ml polybrene (Sigma) in 12-well plates (Sigma). The plates were spun at 1000 × g for 2 hours at 37°C. After spinning, 2 ml fresh D10 media was added to each well and the cells were maintained at 37°C for 24 hours. The cells were then dissociated with TrypLE, suspended in D10, and counted with a TC20 Automated Cell Counter (Biorad). Then, 2,500 cells per well were plated in a black TC-treated 96-well plate with clear well bottoms (Sigma). For each dose, four wells then underwent puromycin selection while four wells continued growth in D10 media. Media was refreshed after 48 hours. After 96 hours, survival of the selected cells in comparison to the unselected cells was measured by CellTiter-Glo (Promega) and used to calculate the lentivirus MOI. The SaCas9 lentivirus was similarly prepared and titered, using blasticidin selection.

### Pairwise library arrayed pilot

As a pilot study, we generated five lentiviral vectors with a shared sgRNA targeting five different target sites, which were mismatched at varied positions to examine specificity. These were synthesized by IDT, cloned into the library lentiviral backbone, and verified by Sanger sequencing. Each construct was individually packaged into lentivirus and titered. HEK293T were transduced by spinfection with the SaCas9 lentivirus and underwent 6 days of blasticidin selection. Then, wells of 3 million cells each were spinfected with the guide-target lentiviruses, in duplicate, and subjected to puromycin selection for 7 days. Cells were harvested every 24 hours. Genomic DNA was extracted and the target sites were sequenced by MiSeq to measure indel rates, using the same computational pipeline as for the pooled screen.

### Pairwise library pooled screening workflow

Starting from a library of 88,692 members and 27% ‘error-free’ reads in the post-tracrRNA plasmid pool; we spinfected 340 million HEK293T cells with the library lentivirus at a low multiplicity of infection (MOI = 0.3) to achieve a desired representation of 300X. 24 hours after spinfection, we began 5 days of puromycin selection. The cells recovered in antibiotic-free D10 media for 1 day. Then, 340 million cells were harvested for the Day 0 timepoint and 820 million cells were spinfected with the SaCas9 lentivirus at an MOI = 0.4 to achieve 1000x representation. They were maintained for 3 days under blasticidin selection. The biological replicate of the screen was performed independently on a different day, starting from the initial library spinfection.

### Genomic DNA extraction, library preparation, and sequencing

Genomic DNA was purified from cell pellets of over 340 million cells with a Quick-gDNA Midiprep kit (Zymo), according to the manufacturer’s instructions. The library-cassette regions from the entire DNA pool were then PCR amplified with primers designed to target the U6 promoter and the constant sequence downstream of the target site. These primers include Illumina sequencing adapters as extensions (P5 on the forward primer and P7 on the reverse primer). There were 12 forward primers and 8 reverse primers, which differed from each other by the length of a ‘stagger’ sequence. The stagger ensures a diversity of base calls in each cycle of sequencing. For each sample, 96 PCRs were performed with: 20 *μ*g gDNA, 25 *μ*l NEBNext Master Mix (NEB), 0.5 *μ*l Q5 polymerase (NEB), 1 *μ*l DMSO, 1 *μ*l MgCl_2_ (25 mM), and 0.5 *μ*M forward and reverse primers, in 50 *μ*l reactions. The thermocycling protocol was: 30 s at 98°C, followed by 18 cycles of 98°C for 10 s, 60°C for 30 s, and 72°C for 30 s, then a final 5 min extension at 72°C. Then, the 96 reactions were pooled. The desired PCR products were then gel extracted and quantified by Qubit. These libraries were subjected to 2×150 bp paired end sequencing on a HiSeq2500 (Genewiz), with 2 lanes per sample.

### Computational analysis

Indel rates were computed with a previously described Python script^23^. Error rates were estimated by comparing the constant regions of the library cassette against reference sequences. The pairwise library pooled screen was analyzed with two pipelines. The first pipeline took the library design file and the Day 0 sequencing reads as inputs and outputted a ‘whitelist’ of validated rBCs, which were associated with error-free library cassettes. The second pipeline took this rBC list and the Day 3 sequencing reads as inputs and outputted the indel rate for each guide-target pair, computed from the subset of reads bearing whitelisted rBCs. This process minimizes false positive indels that may arise from mutations in the target site during the synthesis, cloning, or lentiviral packaging. Off:on-target ratios are an indel rate normalized by the indel rate resulting from the perfectly matched guide of equivalent length. The script was written in Python and run on an Amazon Web Services EC2 instance. Statistical analyses were performed in Graphpad Prism. Two-sided tests were used in all cases.

### GFP reporter of mismatched sgRNA activity

Wild type and mismatched sgRNA were generated by PCR and transfected as amplicons containing U6 promoter, spacer sequence, and tracrRNA scaffold. Singly mismatched sgRNA were designed by swapping T and A, or G and C, at each position. HEK293-GFP were seeded at a density of 100,000 cells/well in 24-well plates. After 24 hours, cells were transfected with 250 ng of gRNA amplicon and 750 ng of wild-type SaCas9 plasmid (pAF003). All transfections were performed in duplicate using Lipofectamine 3000 (Life Technologies). At three and a half days post-transfection, cells had media removed and were washed with 0.5 mL of phosphate-buffered saline (PBS). 200 *μ*l of trypsin was added to the cells and they were incubated at 37 °C, 5 % CO_2_ for 5 min. Trypsinization was halted by adding 0.5 mL of complete media to each well. Cells were collected and transferred to 1.5 mL tubes, spun down at 3000 rpm for 7 minutes, washed with 1.0 mL fluorescence-activated cell sorting (FACS) buffer (PBS with 3 % FBS), spun down again, and resuspended in 200 μl FACS buffer. Cells were then analyzed with a BD Accuri C6 flow cytometer.

### Northern blot

HEK293T cells were transfected with plasmids expressing sgRNA of varying lengths (18, 20, 22, and 24 nt) with a U6 promoter. Samples were co-transfected with a SaCas9 plasmid or the corresponding empty vector. After 3 days, sgRNA were purified from cell pellets with a mirVana kit for small RNA isolation (Thermo Fisher). RNAs were heated to 95 °C for 5 min before loading on 8% denaturing polyacrylamide gels (SequaGel, National Diagnostics). Afterwards, RNA was transferred to a Hybond N+ membrane (GE Healthcare) and crosslinked with Stratagene UV Crosslinker (Stratagene). Probes were labeled with (gamma-^32^P) ATP (PerkinElmer) with T4 polynucleotide kinase (New England Biolabs). After washing, membrane was exposed to phosphor screen for 1 h and scanned with a phosphorimager (Typhoon). The sgRNA expression ratio was quantified based on the intensity of the bands in the image using ImageJ analysis software.

### Model of SaCas9 Specificity

Parameter values for the nonlinear model were derived using Hamiltonian Monte Carlo sampling as implemented in the Rstan package ^24^. The default “No U-Turn Sampler” was used to draw 1500 samples across 8 independent chains, with the first 500 samples in each chain being discarded as part of the “warmup” phase. Convergence of the posterior distributions for the parameters was verified by calculating the R-hat statistic ^25^ and verifying that values for model parameters were < 1.1.

The statistical model is implemented as follows. The indel rate *I*_*j,M*_ observed for a guide *j* targeting a sequence with mismatches *M* is calculated by assuming that the on-target activity of the guide *g*_*j*_ is decreased by additive penalties incurred for each mismatch (ΔΔG_k_) and that the calculated indel rate is subject to independent Gaussian errors ε_j,M_.

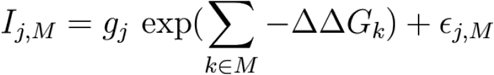

The penalty terms ΔΔG_k_ are restricted to being non-negative and regularized by enforcing a prior distribution that is exponential with mean β = 1. The error terms ε_j,M_ are normally distributed with mean 0 and standard deviation 0.1 (as estimated from comparisons across replicates). w

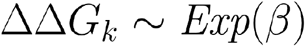

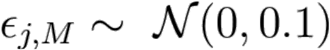

For ease of interpretation, the same model can be reparametrized so that the penalties are multiplicative, as follows:

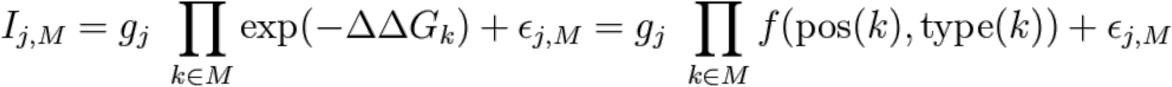

In this case, the penalty on guide activity as a function of the mismatch position and type can be represented as a value between 0 and 1. Parameter values were independently derived for each guide length by separating the data into disjoint training sets, leading to a separate mismatch effect matrix *f*_*L*_ for each value of guide length *L* assayed. The mismatch penalties displayed in the text and figures correspond to the mean of the posterior distribution derived for each of these parameters.

This score fit the observations well for most guides in our dataset, except for guides with very low on-target activity (**Fig. S5B, C**). This may be explained by a low signal-to-noise ratio for these ineffective guides.

## Code availability

The code for the model will be available on the Editas Medicine Github.

## Data availability

The datasets generated during the current study will be available in the NCBI SRA repository.

## Acknowledgements

We would like to thank Team Editas for helpful discussions and support.

## Author Contributions

PDH devised the concept. JT, JSG, and PDH designed the library with input from OOA. JT, NH, and PDH carried out screen. AEF performed GFP knockout validation. XW performed Northern blots. JSG contributed indel detection code. JT and PDH wrote screen analysis code and performed data analysis. LAB designed specificity model. JT, LAB, VM, CW, and PDH interpreted results and wrote the manuscript with help from all authors.

